# Whole-transcriptome RNA editing analysis in single cortical neurons links locus 15q11 with psychiatric illness

**DOI:** 10.1101/840892

**Authors:** Brendan Robert E. Ansell, Simon N. Thomas, Roberto Bonelli, Jacob E. Munro, Saskia Freytag, Melanie Bahlo

## Abstract

**BACKGROUND:** Conversion of adenosine to inosine in RNA by ADAR enzymes occurs at thousands of sites in the human transcriptome, and is essential for healthy brain development. This ‘RNA editing’ process is dysregulated in many neuropsychiatric diseases, but is little understood at the level of individual neurons.

**METHODS:** We quantified RNA editing sites in full-length capture nuclear transcriptomes of 3055 neurons from six cortical regions of a neurotypical post-mortem female donor. Putative editing sites were intersected with sites in bulk human tissue transcriptomes including healthy and neuropsychiatric brain tissue, and sites identified in single nuclei from unrelated brain donors. Differential editing between cell types and cortical regions, and individual sites and genes therein, was quantified using linear models. Associations between gene expression and editing were also tested.

**RESULTS:** We identified 41,930 RNA editing sites with robust read coverage in at least ten neuronal nuclei. Most sites were located within Alu repeats in introns or 3’ UTRs, and approximately 80% were catalogued in published RNA editing databases. We identified 9285 putative novel RNA editing sites, 29% of which were also detectable neuronal transcriptomes from unrelated donors. Inhibitory neurons showed higher overall transcriptome editing than excitatory neurons. Among the strongest correlates of global editing rates were snoRNAs from the SNORD115 and SNORD116 cluster (15q11), known to modulate serotonin receptor processing and to colocalize with ADAR2. We identified 29 genes preferentially edited in excitatory neurons and 44 genes edited more heavily in inhibitory neurons including RBFOX1, its target genes and small nucleolar RNA-associated genes in the autism-associated Prader-Willi locus 15q11. These results provide cell-type and spatial context for 1730 and 910 sites that are also edited in the brains of schizophrenic and autistic patients respectively, and a reference for future studies of RNA editing in single brain cells from these cohorts.

**CONCLUSIONS:** RNA editing, including thousands of previously unreported sites, is robustly detectable in single neuronal nuclei, where gene editing differences are stronger between cell subtypes than between cortical regions. Insufficient editing of ASD-related genes in inhibitory neurons may manifest in the specific perturbation of these cells in autism.

## INTRODUCTION

The extraordinary structural and functional complexity of the human brain arises via multiple mechanisms of genetic regulation. The conversion of adenosine to inosine (A>I) in nascent RNA transcripts in the nucleolus by ADAR1 and ADAR2 enzymes, known as ‘RNA editing’, is the most abundant RNA modification in the primate central nervous system, and confers transcriptomic diversity beyond that encoded in the genome (Slotkin and Nishikura, 2013). RNA editing is essential for healthy brain development and increases with age (Hwang et al., 2016).

Dysregulated editing is implicated in epilepsy (Srivastava et al., 2017), glioblastoma (Silvestris et al., 2019), major depression (Lyddon et al., 2013), autism spectrum disorder (Tran et al., 2019) and schizophrenia (Breen et al., 2019). ADAR1 primarily edits adenosine within repetitive regions; ADAR2 primarily edits non-repetitive regions in the brain, and ADAR3 is a catalytically inactive inhibitor of editing (J. B. Li et al., 2009). ADAR2-null mice die *in utero* and partial knock-out neonatal animals succumb to severe seizures (Higuchi et al., 2000). In humans, mutations in ADAR1 cause skin dyschromatosis (MIM 127400) and Aicardi-Goutieres encephalopathy (MIM 615010) with clinical sub-types including striatal and motor neurodegeneration (Gallo et al., 2017; Livingston et al., 2014). Mislocalization of ADAR was also recently reported in human and mouse models of C9orf72-mediated amyotrophic lateral sclerosis (ALS) (Moore et al., 2019). Common SNPs in the ADAR gene family have also been implicated in numerous diseases including intellectual disability, microcephaly, epilepsy (Tan et al., 2020), hippocampal volume, type II diabetes, and aspects of Alzheimer’s disease and lung cancer (Buniello et al., 2019).

Recently thousands of edited sites were identified in bulk RNA sequencing of human tissues including several brain regions (Tan et al., 2017) as part of the GTEx consortium projects. The most well-characterised edited site is an A>I conversion at exonic nucleotide 2,135 of the glutamate receptor subunit transcript *GLUR2*, which produces a Q>R amino acid substitution. This site is edited in nearly 100% of human *GLUR2* transcripts, and limits the calcium permeability of the resulting ion channel, which is thought to dampen neuronal excitability (Nishikura, 2006). Unlike this ‘gold standard’ *GLUR2* site, most editing sites are located in non-coding regions of the transcriptome, particularly within Alu repeats that form double-stranded RNA on which ADAR enzymes act. Editing of non-coding regions can nevertheless affect protein expression via intron retention, splice site variation and altered translation efficiency (Zhou et al., 2013).

The RNA editing landscape has been described both in the healthy brain and in brains from neurological and neuropsychiatric patients. However this has been performed with bulk RNA sequencing methods, and whilst informative, little is known about the regional and cellular specificity of this process in single cells. Unlike gene expression analysis, RNA editing analysis requires greater sequencing depth and transcript coverage, which is more expensive to generate than widely used 3’ single-cell sequencing protocols. Picardi and colleagues previously profiled editing rates at protein re-coding sites in 268 brain cell nuclei from 7 adult and 3 foetal brains (Picardi et al., 2017b), and found a markedly bimodal distribution of edited allele frequency in contrast to the continuous distribution reported in bulk tissue. This suggests that bulk tissue RNA editing measurements belie highly penetrant editing events restricted to certain cell-types or tissue regions. To better understand RNA editing dynamics in single brain cells, and to detect novel sites that may be subsumed in bulk RNA sequencing, we compared RNA editing in more than 3000 single neurons from six cortical regions of the left hemisphere from an individual donor, previously reported by Lake and colleagues (Lake et al., 2016). This data was generated using the SMART-seq full-length transcript capture platform and is enriched for nuclear RNA, thus allowing deep insight into editing of non-coding and pre-RNAs in cortical neurons of the healthy adult brain. We applied rigorous statistical methods to identify clinical and transcriptomic correlates of RNA editing in this dataset, and report differential site- and gene editing across neuronal sub-types and cortical regions. This work represents the largest and most comprehensive analysis of RNA editing in single cells of any biological system to date.

## RESULTS

### Thousands of novel putative RNA editing sites revealed at single-cell resolution

Independent read processing and unsupervised clustering of 3,127 neuronal nuclei from a single donor, originally assayed by Lake and colleagues (Lake et al., 2016), separated nuclei according to the originally-reported neuronal subtypes (SFigure 1). We discarded 72 nuclei due to either high mitochondrial reads (n = 42), or low library complexity (n = 30) (SFigure 2). We therefore quantified RNA editing signals in 3,055 high-quality neuronal nuclear transcriptomes, supported by 9.17bn uniquely-mapped reads. Excitatory neurons from the superior temporal cortex (Brodmann area 41) were the most abundantly represented cell type in this data set (606 cells; 29.7% of total). The mean ratio of inhibitory to excitatory neurons across each cortical region was 0.285.

Adenosine editing produces an inosine base, which is represented as guanosine in RNA sequencing data. Adenosine-to-guanosine substitution was by far the most common variant detected in nuclear transcriptomes, consistent with a strong ADAR-dependent RNA editing signal relative to variants originating from common genomic SNPs (SFigure 3). By filtering on site coverage, prevalence, genomic context, and previous evidence, we detected 41,930 editing sites. An average of 1,850 candidate editing sites were transcribed per cell (4.4% of total), of which 290 (0.7%) were edited (Figure 1a). When site prevalence was considered, on average each site was transcribed in 138 cells (4.5% of all cells) and edited in 22 (0.7%) (Figure 1b). The distribution of minor (G) allele frequencies within nuclei was strongly bimodal as reported previously (Picardi et al., 2017b) (SFigure 4a). When averaged across cells, the minor allele frequency distribution was positively skewed, and previously uncatalogued ‘novel’ sites showed lower overall editing frequencies (Figure 1c; SFigure 4b). Approximately 80% of editing sites were located within non-overlapping regions of protein-coding genes. The majority of these (20,052; 59%) were located in intronic Alu repeats documented in the REDIportal database of human RNA editing sites (Ramaswami and Li, 2013) (Figure 1d). A further 4,245 sites were exonic, including three-prime untranslated regions (2,958 sites), five-prime UTRs (122) and stop codons (6), in broad agreement with previous genome-wide characterization of RNA editing (J. B. Li et al., 2009) (Figure 1e).

**Figure 1.**
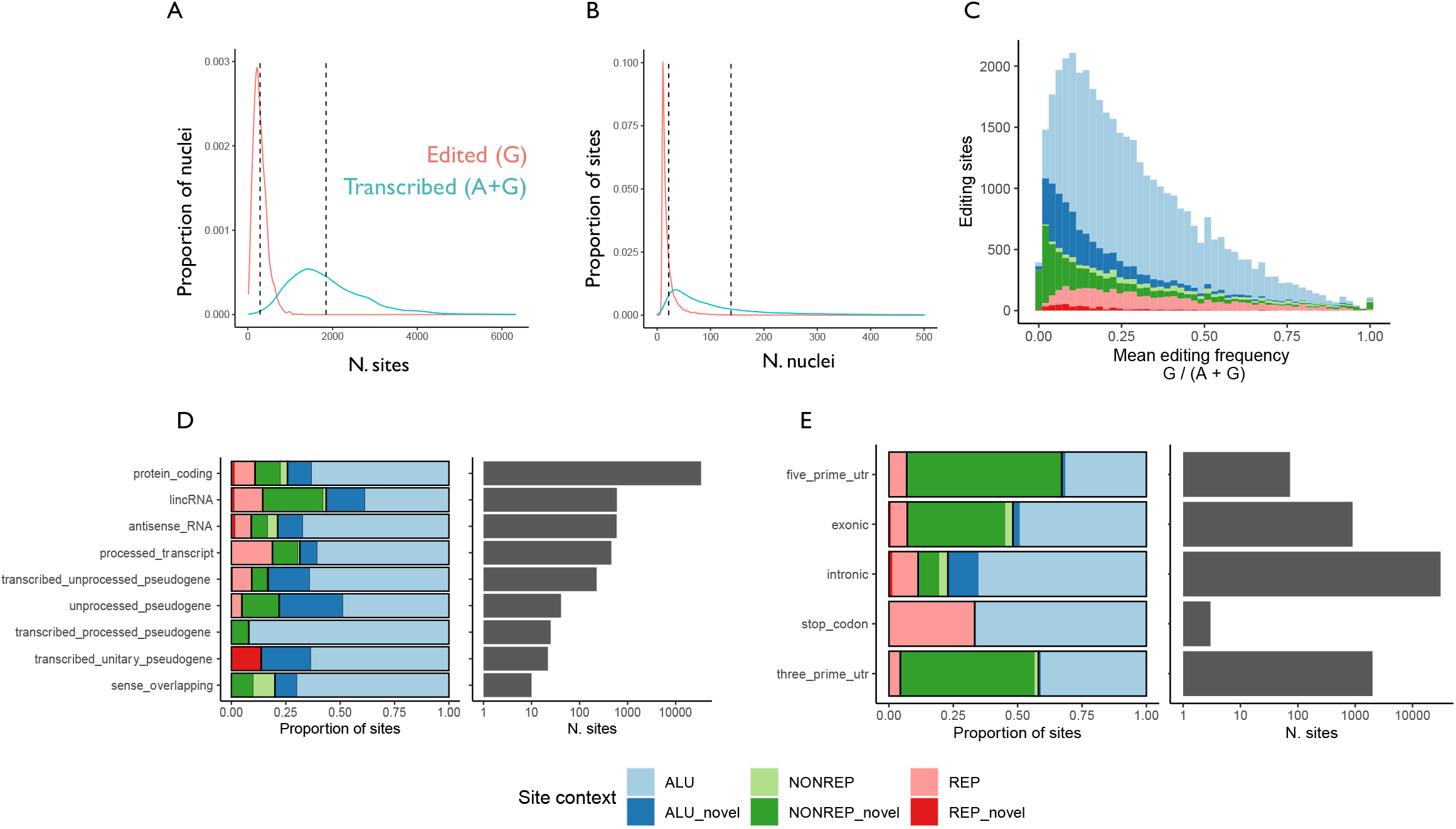
Distribution of 41,930 RNA editing sites across neuronal nuclei, transcriptional frequency, gene biotypes and features. Density of edited and transcribed (including unedited) sites detected A) per nucleus (x axis), and B) across neuronal nuclei (‘nuclei per site’ on x axis). Dashed lines indicate the series mean. C) Distribution of mean minor (edited) allele frequencies for catalogued (light hues) and novel sites (dark hues) located within Alu repeats (blues); non-Alu repeats (reds) and non-repetitive sequence (greens). Individual density plots are provided in SFigure 4b. D) Distribution of sites coloured by site context, across RNA biotypes (y axis). Number of sites is indicated by bars on right panel, on a log scale. E) Distribution of protein-coding sites across gene features (y axis), with number of sites indicated by bars in right panel.

We intersected this data with editing sites documented in 304 bulk RNA sequencing samples from 13 brain regions (36 healthy donors) (Picardi et al., 2017a) (Tan et al., 2017) (STable 1). Some 69.1% of all sites (n = 29,007) were detected in at least one bulk brain sample, of which 30.8% (10,032) were detected in the frontal cortex. A further 3,638 sites were catalogued in other human tissues (Picardi et al., 2017a) (Figure 2a). Of 9,285 newly detected editing sites, 4,547 and 507 sites overlapped Alu repeats and non-Alu repeats respectively (Figure 2b). The remaining 10.1% of sites (n = 4,231) were novel, and located in non-repetitive sequence (Figure 1d, Figure 2b). Across each gene, the strongest predictor of the number of novel non-repetitive sites was the number of novel Alu sites, which in turn was most strongly correlated with the number of documented Alu sites (SFigure 5a). We next intersected our data with differentially edited sites reported based on analysis of bulk brain RNA sequencing of neuropsychiatric patient cohorts, and could provide single-cell level context for 1,730 of 18,071 sites edited in the frontal cortex of schizophrenic (SCZ) patients (9.5% of all SCZ-related sites) (Breen et al., 2019); and 910 of 9,680 sites differentially edited in autism spectrum disorder patients (9.4% of ASD-related sites) (Tran et al., 2019). Taken together, nearly 80% of sites detected in single neuronal nuclei are previously reported in bulk transcriptome sequencing from human tissues, predominantly brain samples from both healthy and disease-affected individuals (Figure 2).

**Figure 2.**
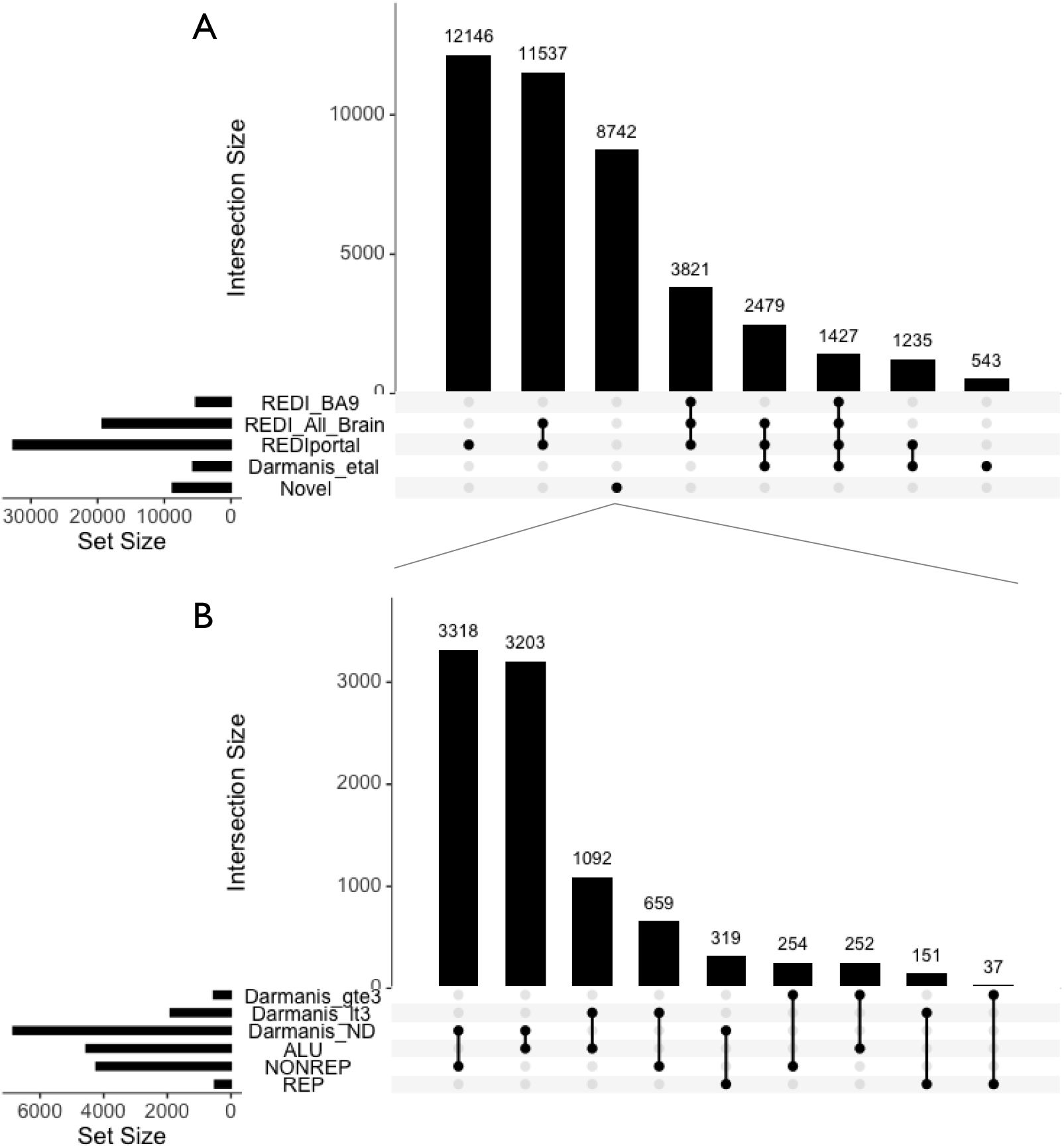
Intersection of RNA editing sites with sites detectable in independently-generated single neurons and bulk tissues. A) Upset plot quantifying editing sites previously documented in the frontal cortex (BA9), or whole brain (≥3 donors respectively), or non-CNS tissues in the REDIportal database (incorporating RADAR and GTEx). Sites detected in at least three of 116 neurons assayed from unrelated individuals (Darmanis et al) are also displayed. B) Intersection of **9,285** previously uncatalogued, novel sites (subset of A) with single neurons published in at least three (gte3) or at least one (lt3) neurons assayed by Darmanis et al; and their editing contexts (Alu, Non-repetitive and repetitive). Horizontal bars indicate set size, and vertical bars indicate size of intersect displayed with linked black points in matrix.

Given the greater sensitivity, cell-type and nuclear-specificity of snucSeq, we sought to validate these findings in the only other publicly available snucSeq from healthy human brain, containing 116 neuronal nuclei from six unrelated donors, published by Darmanis and colleagues. Despite representing only 5% of the number of nuclei assayed by Lake and colleagues, we detected 14% of sites (5,684) from the larger dataset in at least three nuclei, and a further 30.5% of sites (12,781) in at least one nucleus published by (Darmanis et al., 2015) (STable 1). Importantly, 26% of newly detected sites were also identified in this independent snucSeq data (Figure 2b). This demonstrates that RNA editing in single cells is biologically conserved and can be reproducibly detected across unrelated individuals. Further, as much as 20% of editing signals detectable in single-nucleus data may derive from low-penetrance or cell type-restricted modifications that are subsumed by bulk RNA sequencing. All editing sites and associated information for single-nucleus sequencing data reported here are available in supplementary tables, and in an interactive web viewer (**https://shiny.wehi.edu.au/ansell.b/sc_brain_browser**).

### Predicted functional effects and clinical associations of edited sites in single neurons

Functional effects (excluding ‘intron’, ‘up/downstream’ and ‘non-coding’) were predicted for 8.9% of sites (3,736/41,930), with 6.5% of this subset (n = 316) linked to amino acid substitutions. We found edited sites in seven well-established targets of ADAR enzymes that produce modified peptides upon editing, of which four (in *NEIL1, GRIK2, CYFIP2*, and the ‘gold standard’ site in *GLUR2*) had missense mutations which were identical to those previously reported (Nishikura, 2016) (STable 2). All but 12 of the 316 detected missense sites were novel, in non-repetitive exonic regions of 232 genes not currently established as targets of ADAR enzymes. Glutamate- and aspartate-to-glycine substitutions were the most common predicted protein re-coding consequences of editing at these sites (SFigure 6a). We searched for enriched gene ontology molecular function terms among the 232 putative target genes and found both ‘RNA binding’, and ‘adenyl ribonucleotide binding’ among the top 20 most enriched terms— the latter result deriving from missense sites in DEAD box helicases, HSP90 alpha AA1 and AB1, and several calcium transporting ATPases(SFigure 6b).

### Global editing rates differ between neuronal sub-groups and cortical regions

To compare editing rates between neuronal type and cortical region, we calculated a ‘global editing index’ (GEI), taken as the mean minor allele (G) frequency across the sites transcribed in each nucleus (STable 3). Although there was no significant difference in the number of reads mapped to grouped excitatory or inhibitory neurons, a modest effect of library size on GEI was detected, and controlled for in statistical testing. Inhibitory neurons displayed significantly greater editing than excitatory neurons (*p* < 1E-30), which was largely attributable to greater editing in the In6, and lower editing in the Ex3 neuronal subgroups identified by Lake and colleagues (Figure 3c). These trends remained when more conservative (i.e. less sensitive) editing metrics were used, based on Alu sites only (‘Alu editing index’), or Alu sites transcribed in at least 100 cells (SFigures 5b & c). Differences in mean GEI between cortical regions was related to different proportions of neuronal subtypes (Figures 3a & b). Specifically, the visual cortex (BA17) was enriched for neurons of sub-group Ex3, and showed both the lowest proportion of inhibitory neurons (18%), and the lowest mean GEI (see STables 4 and 5 for linear modelling results). Similarly the superior temporal cortex (BA41) was enriched for group Ex1 neurons, and showed a lower mean GEI than the frontal cortex (BA8 and BA10) in which inhibitory neurons (In1, In6 and In8) were more abundant.

**Figure 3.**
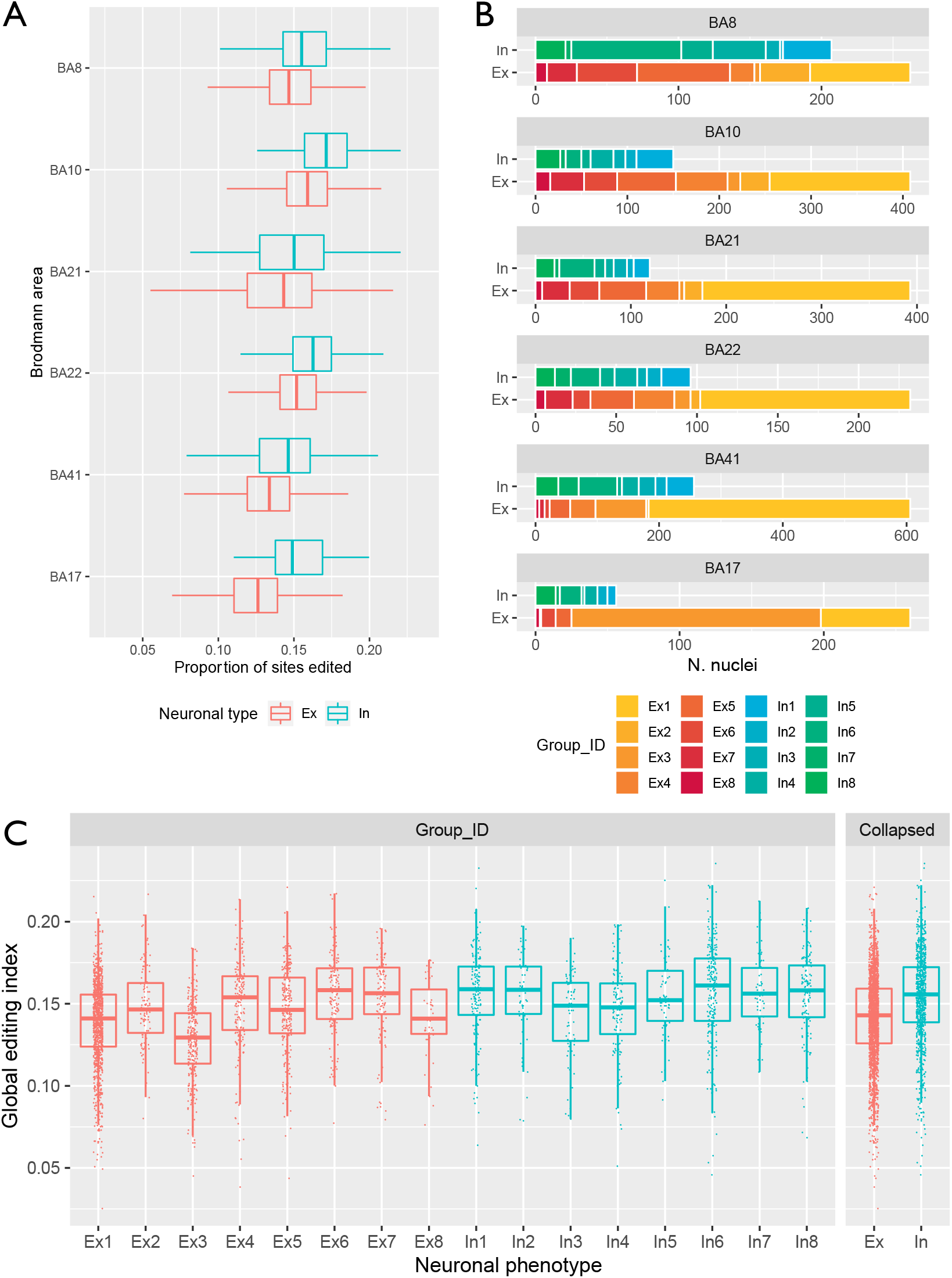
Editing differences between cortical regions and neuronal subtypes. A) Distribution of global editing index (GEI; i.e., mean minor (G) allele frequency for sites in each cell) across six cortical Brodmann areas (BA), for inhibitory (teal) and excitatory (red) neurons. All groups except for Ex neurons in BA10 and In neurons in BA22 are show significantly different GEI compared to In neurons of BA8. B) Abundance of neuronal sub-groups between cortical regions, as defined by (Lake et al., 2016). Note different x axis scales. C) Global editing index in neuronal sub-groups, and when collapsed into gross neuronal phenotypes. All sub-groups are significantly different to Ex1 except for Ex8 and In3. In: inhibitory neuron; Ex: excitatory neuron.

### Differential gene editing

To further interrogate RNA editing at the level of individual sites and genes, we aggregated single-cell editing signals into ‘pseudo-bulk’ RNA samples and applied a differential editing framework which has been previously employed to quantify differential DNA methylation (Chen et al., 2017). Briefly, this method uses multivariate generalised linear models to identify significant differences in edited (alternative) and unedited (reference) allele counts between grouped cell transcriptomes (see Methods). We compared editing between excitatory and inhibitory neurons by creating pseudo-bulk RNA samples (collapsing RNA-editing counts across cells occupying 16 neuronal sub-groups identified by Lake and colleagues), and correcting for cortical region of origin. Whereas no individual site was significantly differentially edited (dEd) after correction for multiple testing, this framework was amenable to rotation testing used in gene set enrichment analysis (Wu et al., 2010), which revealed 73 genes that were enriched with edited nucleotides in one neuronal sub-type (FDR < 0.1; STable 6). Among 29 genes preferentially edited in excitatory neurons were the putative glutamate transporter gene *SLC38A6*, and the neuronal growth and migration factors *PKN2* and *FAM19A2.* Editing in transcriptional regulators was particularly notable, in *RFX3, POU2F2, CHD6, CDK17, SSBP2, HAT1* and *SLC44A5* (Figure 4a). Lastly among transcripts preferentially edited in excitatory neurons were pro-survival proteins (*LATS1, BMPR1A, BMPR2* and *WDR26*), cell cycle and division proteins (*MNAT1, MZT2A*), and nervous system development proteins (*EXOC6B, COL4A3BP*).

**Figure 4.**
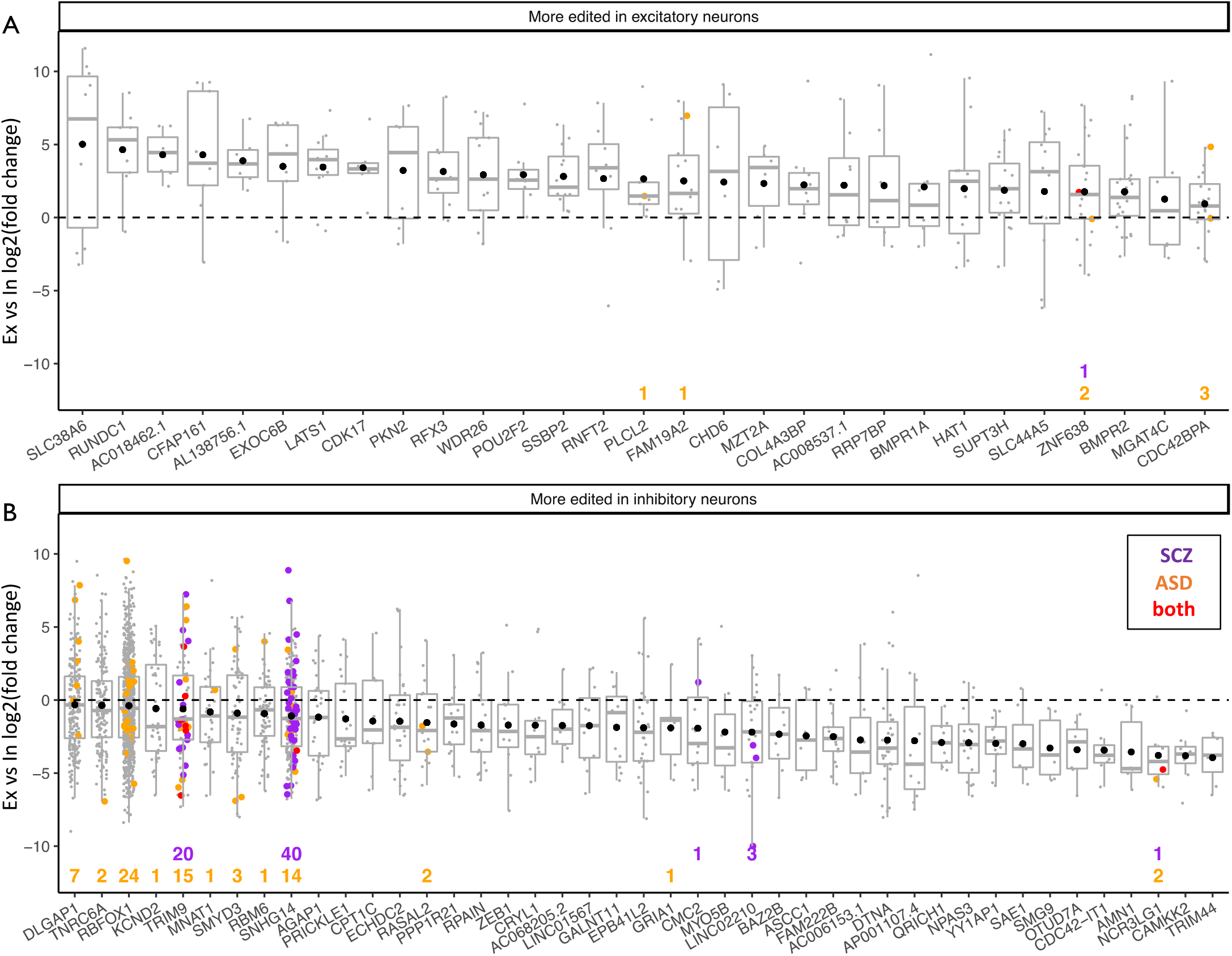
Differential gene editing between excitatory and inhibitory neurons. Genes with preferential editing in excitatory neurons (A) or inhibitory neurons (B). Each point represents log2 fold-change (y axis) in an edited site in a gene (x axis). Box plots indicate distribution of middle 50% of editing fold-changes (box boundaries) and median fold-change (middle line). Black points represent the mean fold-change in editing per gene. Sites differentially edited in cohorts of individuals with schizophrenia (SCZ; purple) or autism spectrum disorder (ASD; orange) in both cohorts (red) are coloured accordingly. The relevant number of such sites per gene is displayed numerically at the bottom of the panel.

The remaining 44 dEd genes were more edited in inhibitory than excitatory neurons. Included in this set was RBFOX1, a splicing regulator, and a further eight binding target genes of RBFOX1 (Weyn-Vanhentenryck et al., 2014). The small nucleolar host gene *SNHG14* was also preferentially edited in inhibitory neurons, as were transcriptional regulators and RNA-binding proteins, represented by *YY1AP1, ZEB1, NPAS3* and the RNA-interference enzyme *TNRC6A*. Preferential editing of several post-translational modification enzymes was notable, including the sumoylation enzyme *SAE1*, histone methyltransferase *SMYD3*, and the de-/ubiquitylation enzymes *TRIM9, TRIM44, OTUD7A, CDC42* and *BAZ2B.* Interestingly, editing at four of five candidate sites in the AMPA glutamate receptor subunit 1, which oligomerizes with GRIA2, was more prevalent in inhibitory neurons. The excitatory neuronal marker gene *CDKK2* also showed a strong editing bias in these cells. Lastly, preferential editing in inhibitory neurons was identified in genes involved in mitochondrial metabolism: carnitine palmitoyltransferase 1C, and metallochaperone *CMC2*; as well as genes related to neurogenesis and synaptic function (*ASCC1*, *DTNA*, *DLGAP* and *EPB41L2*).

A similar analysis of differential editing between cortical regions, correcting for cell type, identified 99 dEd sites across 90 genes. Whereas no gene was significantly enriched with edited sites in any region, the majority of dEd sites were identified in the frontal cortex (BA8: 48 sites; BA10: 24 sites), with 14 dEd sites identified in the temporal cortex (BA22), and fewer than ten in other cortical areas. Three individually significant dEd sites were identified in RBFOX1 and RIMS2 respectively, with two sites in *DNAJC11*, *EPN2*, *PEAK1* and *TPST1* (STable 7). Taken together these results suggest that major editing differences generally reflect neuronal phenotype rather than cortical region of origin.

### Intersection of differentially edited sites in neuropsychiatric illness with neuronal subtypes

Hundreds of differentially edited sites are reported in the brains of individuals with schizophrenia (Breen et al., 2019) and autism spectrum disorder (ASD) (Tran et al., 2019) relative to healthy controls. We tested whether genes that were dEd between inhibitory and excitatory neuronal nuclei were enriched with sites related to these conditions. Indeed, 73 sites dEd in ASD were localized within 12 genes that were more edited in inhibitory than excitatory neurons. In the same manner, 60 of 65 sites dEd in schizophrenia were concentrated in the *TRIM9* and *SNHG14* genes. Conversely, only seven dEd sites in ASD and one in schizophrenia were identified among genes displaying increased editing in excitatory neurons (Figure 4a). Although dEd sites associated with these conditions were 2.6 times more likely to be present in genes dEd in inhibitory neurons, this was not significant by Fisher’s exact test (*p* = 0.16) and will require further analysis in an independent data set.

### Small nucleolar RNA expression is a marker of nuclear RNA editing

Expression of ADAR family enzymes is a modest predictor of editing activity across human tissues (Roth et al., 2019; Tan et al., 2017), which has prompted a search for other modulators of editing (Quinones-Valdez et al., 2019; Tan et al., 2017). To identify such molecules we modeled GEI as a function of gene expression, controlling for neuronal type. A total of 170 genes were significantly associated with GEI, with absolute effect sizes greater than 0.01 (FDR < 0.05)(STable 8). As these genes were among those most consistently expressed across cells (mean n. cells per gene = 2,250 and 687 for significant- and non-significant genes respectively), we down-sampled the data to a maximum of 1,500 cells per gene, and found that all results remained significant. Interestingly, expression of ADAR family genes was not associated with GEI at the single-nucleus level. By contrast, 47 small nucleolar RNA genes transcribed from the locus 15q11 showed strong positive correlations with GEI (Figure 5a). This result was due to independent contributions from both neuronal subtypes, and was robust to the removal of a large cluster of edited sites in the precursor transcript *SNHG14* (Figure 5b). Another four snoRNAs from the *SNORD3* cluster on chromosome 17 were detected together with *SNORD114* (chromosome 14) and H/ACA snoRNAs transcribed from chromosomes 1, 3 and 20. Other non-coding RNAs including three small nuclear RNAs and eight ribosomal RNAs were also positively associated with GEI. Only two protein-coding transcripts in this group were identified: the mitochondrial carnitine transporter *CACT*, and the lipid transport protein apolipoprotein A.

**Figure 5.**
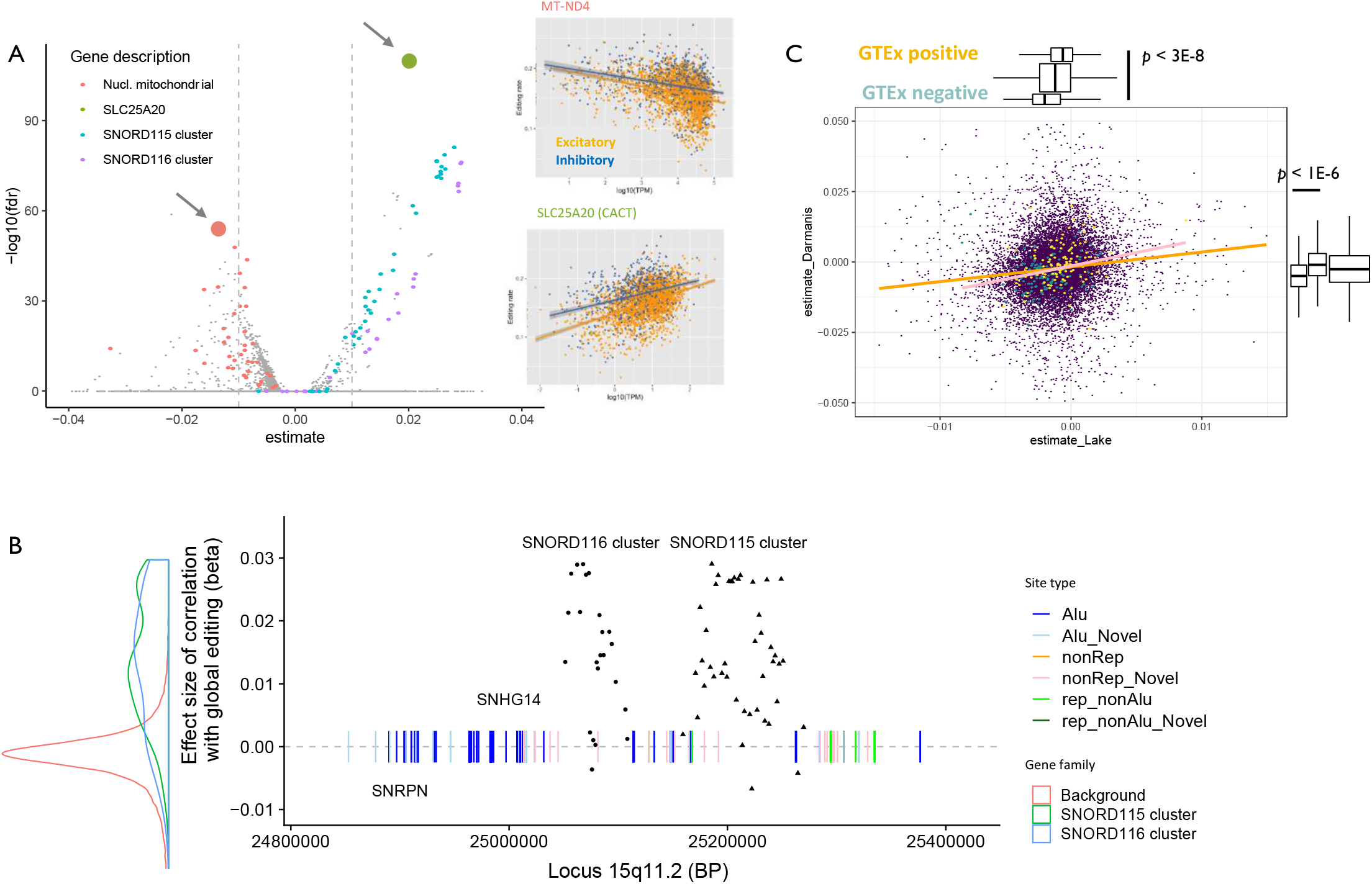
Associations between gene expression and global RNA editing rates. A) Associations between log10(gene expression) and the cell-wise global editing index are represented as effect size (x axis) and transformed FDR-adjusted p value (y axis; higher values are more significant). Inset displays correlation scatter plots between global editing (y axis) index and log10 gene expression (x axis) for neuronal nuclei expressing representative significant results: mitochondrial NADH oxidase 4, and CACT (marked with arrows). B) Expression SNORD115 and 116 snoRNAs in locus 15q11 correlates strongly with global RNA editing. Genomic location is displayed in base pair units on the x axis; effect size of correlation with editing is displayed on the y axis for SNRPN, SNHG14 (grey arrows) and SNORD cluster genes (black points). The distribution of correlations between the SNORD clusters and editing is displayed at left relative to all other genes (background), and scaled to the y axis. Edited sites are represented by vertical lines, and coloured according to genomic context. C) Correlation between gene expression-editing associations in two single nucleus datasets. Each purple point represents a gene. Associations between log expression and global editing in the Lake et dataset (x axis) are displayed relative to results from Darmanis et al dataset (y axis). The orange line indicates fitted correlation between datasets; the pink line indicates correlation between a subset of genes (yellow and green points) identified to have RNA editing effects in GTEx data. Marginal box plots display significant separation of positive (yellow) and negative (green) results from GTEx in both snucRNA datasets.

Transcripts whose expression was negatively associated with editing, related to trans-synaptic signaling (calmodulins 1 and 2, microtubule associated proteins *MAP1A* and *1B*, *SNAP25*, and syntaxin binding protein 1), and metallopeptidase protein-coding genes (*DPEP1, MMP10* and *TRABD2B)—* some with roles in cellular differentiation (ADAM metallopeptidase TS20). Greater expression of coagulation factor X and leucine-rich repeat protein 15 was also negatively associated with GEI, as was expression of seven long non-coding RNAs. Also in this group were the Alu repeat-containing transcripts *7SL* and olfactory receptor *OR52A5*, and genes related to neurogenesis (chimerin 1 and olfactomedin). Transcripts encoding proteins with transcription factor activity (*PAX6*, *IRX2*) and several related to neurotransmission (*LYPD4*, tryptophan 5-monooxygenase, the active calcium transporter *ATPB2* and potassium channel protein *KCNJ15*), were also negatively associated with GEI. Lastly heat shock protein 90 and mitochondrial NADH oxidase transcripts showed a negative relationship with editing. When the effect-size threshold was omitted, we observed enrichment of nuclear-encoded mitochondrial genes among the negative transcriptional correlates of editing (Figure 5a).

To test the consistency of these findings across adult neurons, we performed the same analysis on 116 neuronal nuclear transcriptomes assayed by (Darmanis et al., 2015) (STable 9). Whereas ADAR enzymes were again not significantly associated with editing, the calmodulins 1 and 2, SNAP25, beta actin, MAP1B and protein phosphatase 3CA transcripts showed negative associations in both datasets. There were no common genes positively-associated with editing between the two datasets, which may be due in part to significant differences in the classes of assayed RNA biotypes (SFigure 8). Nevertheless, the correlation between editing associations (beta values) was positive and significant (R = 0.11, *p* < 1e-30; Figure 5c). Furthermore, genes with positive editing associations in bulk RNAseq from human tissues (Tan et al., 2017), showed similar positive trends in results from both single-nucleus datasets. Conversely, genes with negative editing associations in GTEx tissues showed congruent negative associations in single cell data (R = 0.26, *p* < 1E-4), and were significantly segregated from positively-associated genes (Fig 5c marginal boxplots). Finally, as raw SMART-seq data from human brain is exceedingly rare, we investigated the association between 15q11 snoRNA expression and the GEI in 20 bulk brain RNA libraries from five adult donors, generated using a ribosomal RNA depletion protocol wherein snoRNAs were readily detectable (Hwang et al; SFigure 7). After correcting for brain donor ID and estimated cell-type proportions, the mean expression of the SNORD116 cluster, but not SNORD115, was positively associated with global tissue editing, driven by higher global editing and expression of SNORD116 in the cerebellum (SFigure 8). Taken together, these results suggest extensive involvement of small nucleolar and other non-coding RNAs in RNA editing, and a spectrum of transcriptional associations with editing that is broadly conserved in single neurons and bulk brain tissue.

## DISCUSSION

RNA editing is ubiquitous in the human central nervous system, and aberrant editing is implicated in numerous neurological/neuropsychiatric diseases, cancer and immune disorders. We undertook the first high-resolution exploratory investigation of this process in single neuronal nuclei assayed across 6 cortical regions from a healthy adult brain donor, and found extensive agreement with previously reported edited sites in healthy and disease-affected brain tissue. Further, the abundance of nuclear/nucleolar pre-RNA species in this dataset profiled with full-length capture technology, facilitated unparalleled insight into editing activity across the transcriptome occurring in physical proximity to brain-specific ADAR enzymes (Gallo et al., 2017). By applying a sensitive linear modeling approach to detect differential site- and gene editing across cell types and cortical regions, we demonstrated that RNA editing analysis is feasible in snucSeq, and informative for investigating disease-relevant sites in the context of specific neuronal cell types. We further identified editing differences in disease-related genes between neuronal subtypes, and a strong association between snoRNA expression and transcriptome-wide RNA editing.

This work was made possible by publicly available sequencing data and curated databases. The novel sites identified here require further validation and functional studies, and will constitute the first publicly available cell-type specific editing reference database (available at **shiny.wehi.edu.au/ansell.b/sc_brain_browser**). The main limitations of this study lie in the lack of RNA stranding information in the SMART-seq data, and the lack of reference genomes for brain donors— both of which hamper our ability to disambiguate editing sites from genomic SNPs. To address these concerns we took stringent measures including: i) removing common genomic SNPs from the results, ii) removing sites derived from transcripts that overlap on opposing strands; and retaining only those sites iii) detected within genes on the cognate strandiv) covered by at least five reads, and vi) detected in at least ten nuclei. The latter filter should eliminate PCR and other random artefacts because SMART-seq library preparation is independent for each nucleus. Although brain SMART snuc-RNAseq from healthy donors is exceedingly rare in public databases, we also called RNA editing sites *de novo* in the only other publicly available dataset of this kind (Darmanis et al., 2015), and assessed the reproducibility of our findings, achieving good correlation, allowing for the smaller sample size of the replication dataset. Lastly, in lieu of additional publicly available single-nucleus neuronal datasets, we validated key findings relating to non-coding snoRNAs using ribo-depleted bulk brain RNA libraries after cell-type deconvolution.

The proportionate distribution of edited sites across RNA motifs (Alu and non-Alu repeats) and genic features (mostly intronic or 3’ UTR) broadly agrees with the original characterization of editing in the human transcriptome (J. B. Li et al., 2009). Excitingly, thousands of putative novel editing sites were detectable in the present data, thanks to the extensive coverage of the transcriptome at single cell resolution. That a substantial proportion (58%) of 9,285 newly identified sites occurred within canonical Alu/repetitive non-Alu contexts, and 26% were detectable in independent sequencing of neurons from unrelated individuals, suggests that this data type contains novel and reproducible insights which are not detectable in bulk RNA sequencing. Indeed, the cellular heterogeneity of RNA editing is a striking result in the present study.

Whereas previous studies of bulk RNA portray editing as a pervasive low-frequency phenomenon (Tan et al., 2017), our results suggest that this is, at least in part, an artefact of high frequency editing activity that is restricted to subsets of cells, and thus masked by ensemble averaging (SFigure 4). Accordingly, a fundamental re-evaluation of our understanding of this process is warranted, and routine integration of editing and expression data in single cell studies has great potential to inform the structural and functional heterogeneity of the brain. As the present data is enriched for nuclear pre-RNA, it likely predominantly reflects editing activity of ADAR2, which is spatially restricted to the nucleus and predominantly expressed in neurons. Conversely, highly prevalent but low-penetrance editing, and sites edited in cytosolic RNA, will be under-represented in the current data.

### Site-specific editing

Our study was able to quantify editing sites across cortical regions and neuronal subtypes with high resolution, as well as investigate the transcriptomic underpinnings of neurological and psychiatric illness in the context of cell-type and brain region.

We found that a number of transcripts encoding putative novel missense sites are involved in adenosine binding. Editing-based feedback loops in editing-related enzymes are reported in model systems (Feng et al., 2006), and may be more extensive than previously thought. Beyond adenosine binding proteins, the presence of putative recoding sites in synaptic transmission-related proteins, as well as solute carrier family members and neurotransmitter receptors, suggests that editing-based peptide modification modulates both the sensitivity of neurons to stimuli, and the responsivity at the synapse.

### Differential editing between cell types and cortical regions

Comparison of the global editing index (‘GEI’) between cell-types and cortical regions was facilitated by averaging the editing frequencies of transcribed sites in each cell. Both cell types in the frontal cortex exhibited a higher GEI than other regions. Interestingly, Brodmann areas 17 and 41 house the primary visual and auditory cortices, respectively, and exhibited lower global editing rates. If higher editing rates are assumed to dampen neuronal activity in both inhibitory and excitatory cell types, then these regional differences may support the prevailing understanding of the functional organization of the cortex, wherein sensory regions provide strong excitatory input (corresponding to reduced editing); and the frontal cortex is primarily inhibitory, acting to increase the salience of selected perceptual inputs (Hu et al., 2013).

Using a novel pseudo-bulk framework for RNA editing analysis that has been applied previously to quantify differential DNA methylation (Chen et al., 2017), permitted detection of differences in site-wise and gene-wise editing with great sensitivity. This approach benefits from the ability to apportion biological and technical variation under a negative binomial model of raw count data. Interestingly, whereas differences between cortical regions manifested in differentially edited sites alone, editing differences between cell-types were significant in aggregate at the gene level, but not at the site level suggesting that the volume rather than identity of edited sites across transcripts may distinguish different cell types. Differentially edited genes in excitatory neurons tended to contain relatively small numbers of edited sites showing large fold-changes. This is consistent with the lower global editing rates observed in excitatory cells. Prominent editing of a putative glutamate transporter, *SLC38A6*, is notable given the high requirement for glutamate— the primary excitatory neurotransmitter, in these cells. Nearly a quarter of genes preferentially edited in excitatory neurons encode transcription factors or histone modification enzymes, pointing to potential feedback between editing and transcription. Whereas *RFX3*, *POU2F2*, *CHD6* and *CDK17* or their orthologues have defined roles in the brain, the roles of *SSBP2*, *CDK17* and *HAT1*, and editing therein, newly identified in this study, require further investigation. In addition, heightened editing in excitatory neurons of transcripts involved in self-renewal of neural progenitors (*LATS1*) and pro-survival proteins (*WDR26, BMPRs 1A* and *2*), suggests that a primary role of editing in these cells could be to contribute a homeostatic signal. This is supported by negative associations between editing and expression of both nuclear mitochondrial genes, and four cell-type markers of the Ex1 subgroup. It may be that pro-survival transcripts act as sensors of excessive mitochondrial respiration, which impinges on RNA editing and viability in excitatory neurons. We stress however that the negative association between mitochondrial gene expression and editing is also seen in bulk tissue sequencing (Tan et al., 2017), and is unlikely to reflect apoptosis in this case, as any nuclear libraries containing more than 15% mitochondrial RNA were removed during pre-processing. Complex interactions between mitochondrial metabolism and editing are also suggested by the strong positive association between editing and expression of the carnitine shuttle protein, *CACT*, required for fatty acid oxidation. Dysregulated editing of carnitine pathway genes was recently reported in ALS (Moore et al., 2019), and now warrants further molecular investigation.

Several transcripts assayed in inhibitory neurons exhibited high-density editing, namely *RBFOX1*, *TNRC6A*, *DLGAP1*, *TRIM9* and *SNHG14*. These genes also encompassed vastly more sites reported as differentially edited in neuropsychiatric patient cohorts, most notably autism spectrum disorder (ASD). Sites linked to schizophrenia were largely restricted to the ubiquitin E3 ligase *TRIM9*, and *SNHG14*. The former protein may promote editing in inhibitory neurons by dampening the function of *PAX6*, whose expression is negatively associated with editing in our data. Indeed, cross-talk between post-transcriptional and post-translational modification systems in these cells is also suggested by editing in transcripts central to synaptic formation including de/ubiquitylation enzymes (*TRIM* family and *OTUD7A*), *DLGAP1* and *DTNA*.

Increased editing of the ASD ‘hub’ gene *RBFOX1* (Voineagu et al., 2011), its target genes (Weyn-Vanhentenryck et al., 2014), and other genes implicated in ASD (Parikshak et al., 2013), aligns with previous work implicating dysfunction of inhibitory interneurons specifically in this disorder (Lunden et al., 2019). RBFOX family genes are implicated in excising C/D box small nucleolar RNAs from their host gene *SNHG14* (Coulson et al., 2018), which was also heavily edited in inhibitory neurons. Genomic deletion of the SNORD115 or 116 cluster is sufficient to cause Prader-Willi syndrome (Bieth et al., 2015), which features autistic traits (Descheemaeker et al., 2006). Although *SNORD115* RNAs are known to direct editing and splicing of *5HT2C* serotonin receptors (Bratkovič et al., 2018), we identified a striking positive association between expression of *SNORD115/116* RNAs and editing across the entire transcriptome, which has not been observed previously. This association may have been revealed due to the relative enrichment of nucleolar pre-RNA in this dataset. Small RNAs are rather poorly represented in the Darmanis et al snucSeq data (Darmanis et al., 2015), and especially so in poly-A enriched bulk brain RNA sequencing.

We hypothesize that locus 15q11 snoRNAs may recruit ADAR enzymes or stabilize their interactions with substrates in the nucleolus. Alternatively, these molecules may compete with ADARs for RNA substrates to fine tune editing. Indeed, the abundance of genes whose expression is negatively correlated with editing in this study, and findings associating many more RNA binding proteins with negative rather than positive effects on editing (Quinones-Valdez et al., 2019), suggest that this epi-transcriptomic process may be constitutively active and subject to negative regulation. Together these findings indicate that the 15q11 region is a nexus of RNA editing-related activity whose transcription is a proxy for global RNA editing rates. The extensive differential editing reported in 15q11 in this study between neuronal sub-types and cortical regions; and in two prominent neuropsychiatric disorders, makes this region a compelling subject for further investigation. It is also intriguing to consider the potential involvement of RNA editing dysregulation in neurodevelopmental conditions associated with structural variation in this locus (Jønch et al., 2019; Ulfarsson et al., 2017).

The lack of significant correlation between ADAR family gene expression and editing in either single-nucleus dataset is surprising but not unprecedented given that expression of the neuronal nucleus-restricted ADAR2 explains only 3% of variation in editing of repetitive sites including Alus (Gallo et al., 2017; Tan et al., 2017), which constitute the majority of the present data set. We also emphasize that the relatively small effect sizes for associations between gene expression and editing found here, are of a similar scale to those reported in analyses of bulk RNA that produced novel experimentally-validated modulators of editing (Tan et al., 2017). Despite this, the overall agreement in the sign and magnitude of transcription-editing associations across sequencing technologies in independent studies suggests that a complex network of interacting partners including non-coding RNAs may affect RNA editing, and await experimental validation.

In conclusion, we integrated RNA editing signals in individual neurons with known editing sites in the healthy and diseased brain. Thousands of newly reported editing sites share broad characteristics of documented sites and replicate in independent single-cell data, but are likely subsumed by ensemble averaging in bulk tissues. Sensitive detection of differential site- and gene-editing revealed concentration of ASD-related gene editing in inhibitory neurons. Expression of the ASD-associated Prader-Willi locus snoRNAs mark global editing activity, and associations between editing and transcription are conserved across studies. This work adds exciting new dimensions to our understanding of post-transcriptional regulation in healthy adult cortical neurons, and a comprehensive reference for forthcoming investigation of editing in single-cell samples from neuropsychiatric patient cohorts.

## DECLARATIONS

The authors declare no competing interests.

## MATERIALS and METHODS

### RNA sequencing data and reference databases

Raw data from full-length (SMART-seq) sequencing of single brain cell nuclei was downloaded from dbGaP (study accession phs000834; project ID 20576) (Lake et al., 2016). The neuronal sub-type annotations assigned by the study authors based on gene expression profiles (8 excitatory and 8 inhibitory neuronal sub-groups) were also downloaded. Single nucleus SMART-seq data for 116 neurons from the anterior temporal lobe of four adult donors (GSE67835) (Darmanis et al., 2015), and ribo-depleted bulk RNA-seq libraries from adult human brain tissue (SRP076099) (Hwang et al., 2016), were downloaded from the Gene Expression Omnibus using the NCBI SRA toolkit. Metadata including the cell type and cortical area of cell origin were accessed from the sequence read archive and/or relevant supplementary materials (STables 1 and 2). The REDIportal database (Picardi et al., 2017a) which incorporates the RADAR (Ramaswami and Li, 2013) and GTEx human tissue RNA editing databases (Tan et al., 2017) was downloaded from http://srv00.recas.ba.infn.it/atlas/download.html. Differentially edited sites reported in neuropsychiatric brain analyses (Breen et al., 2019; Tran et al., 2019), were downloaded, and where required sites were transposed from Hg19 to Hg38 using liftOver software (Hinrichs et al., 2006). Genomic ranges for Alu and non-Alu repeats in Hg38 (RepeatMasker track) were downloaded from the UCSC Table Browser (Kent et al., 2002); common human single nucleotide polymorphisms were downloaded from dbSNP (GRCh38p7; build 151) (Sherry et al., 2001). Genes with known amino acid re-coding sites, those implicated in autism spectrum disorder (SFARI database), and targets of RBFOX splicing were retrieved from tables provided in the relevant literature (Abrahams et al., 2013; Nishikura, 2016; Weyn-Vanhentenryck et al., 2014). Variants associated with clinical conditions were accessed via the ClinVar database (Landrum et al., 2018).

### RNA sequence processing and gene expression quantification

RNA sequencing reads were mapped to the human reference genome (GRCh38.91) using the STAR aligner (Dobin et al., 2012) in two-pass mode with the splice junction database overhang maximized for the respective read lengths (47 for phs000834; 74 for GSE67835; 99 for SRP076099). Gene expression was quantified using featureCounts with fractional assignment of multi-mapping reads (Liao et al., 2014). Single-nucleus SMART-seq datasets were further filtered to remove samples with abundant mitochondrially-encoded RNAs (likely indicating apoptosis), low library complexity and low coverage. Count libraries were normalized for size and complexity, and clustered using scran (Lun et al., 2016) and scater (McCarthy et al., 2017). For cluster analysis of the phs000834 data, nuclei were coloured according to the neuronal phenotype and cortical region designated by Lake and colleagues (Lake et al., 2016).

### RNA editing detection and filtering

We called single nucleotide polymorphisms (SNPs) in the nuclear transcriptomes using the GATK best-practices pipeline (Van der Auwera et al., 2013) for RNA-seq data. This involved read deduplication, splitting, base quality score recalibration, and variant calling with HaplotypeCaller. No call confidence filter was applied at this step in order to retain all potential variants. Variant call format files were converted to GDS format using the R SeqArray package (Zheng et al., 2017), and combined with common genomic SNPs, and REDI sites. For phs000834, A>G and T>C sites covered by at least 5 reads, with a minor allele count of at least 2, and detected in at least 10 high-quality (i.e., non-apoptotic, high transcriptional complexity) libraries, were retained.

Common genomic A>G and T>C SNPs were extracted for use downstream. As SMART-seq data is unstranded, categorically determining the originally edited strand is not possible. Therefore to enrich the dataset for true editing sites, after removing common genomic SNPs, we imposed further filtering as follows. Sites were only retained if they were: a) present in previously published RNA editing database, and/or b) located within non-overlapping regions of an ensemble feature, and on the cognate strand (i.e. A>G sites within a feature on the 5’ strand; T>C on the 3’ strand). The bedtools intersect module was used to locate previously unreported sites within genomic repeats (RepeatMasker); and to identify the genic feature (e.g. start codon, exon, intron, 3’ UTR etc) occupied by sites within protein-coding genes (Quinlan and Hall, 2010). Sites with insufficient coverage were distinguished from transcribed, un-edited sites using the samtools depth module (H. Li et al., 2009). The Variant Effect Predictor (McLaren et al., 2016) was used to predict the molecular consequences of RNA editing sites. Only predicted effects other than ‘up/down-stream gene variants’, ‘intergenic’ and ‘intronic variants’ were reported. For GSE67835 the same site detection and filtering procedure was followed except a lower prevalence threshold (at least three nuclei) was applied given the smaller number of neurons under consideration.

For strand-specific ribo-depleted bulk RNAseq data, editing sites were identified in de-duplicated mapped reads using JACUSA software in strand-specific mode. The JacusaHelper R package was used to import variant calls with test statistic values ≥ 1.56 and total read coverage ≥ 10. The minimal alternate (i.e. edited) allele depth was set to 3. Common genomic SNPs were discarded and only A>G variants detected in at least 3 of 20 samples from 5 individuals, were retained for further analysis. Intersection of putative editing sites with genomic ranges databases, and total read depth quantification for all high-confidence sites in all samples, was performed as for single-nucleus data.

### Statistical analysis

To assess the replication of editing sites detected in this and other studies, candidate sites that survived filtering were intersected with sites reported in bulk RNA sequencing datasets and single neuronal nuclei (Darmanis et al., 2015) (GSE67835) detailed above.

Differential editing was assessed at the level of whole nuclei, aggregated over all transcribed candidate editing sites; and between genes using a pseudo-bulk framework. A nucleus-level ‘global editing index’ (GEI) was quantified as the mean minor allele (G) frequency for candidate editing sites in each cell cell. This metric was compared with an ‘Alu editing index’ (AEI), composed of Alu sites only; and a more stringent AEI comprising Alu sites transcribed in at least 100 cells. Edited sites were subsequently grouped by transcriptomic context (Alu repeats, repetitive non-Alu sequence, and non-repetitive sequence) to determine the contribution of each to global editing.

Given the sparse coverage in single-nucleus RNA sequencing, in order to robustly detect differential gene-level editing between neuronal subtypes controlling for cortical region, we adapted a counts-based framework to test for differential editing. Specifically, pseudoreplicates were created by summing counts of reference (unedited) and alternate (edited) alleles at sites with count variances > 0.2, across neurons in phenotypically similar sub-groups as defined by Lake and colleagues (**STable 1)**. This allowed testing differential site-wise editing between gross neuronal subtypes whilst controlling for cortical region, using a negative binomial model of gene expression in the limma R package, as has been successfully applied for differential methylation analysis (Chen et al., 2017; Ritchie et al., 2015). In contrast to ratiometric testing approaches, this method takes site coverage into account, thereby minimizing the probability that lowly expressed, highly variable sites are called as significant. Differential editing at the whole-gene level was identified by testing for enrichment of sites with consistent directional fold changes in editing between neuronal sub-types or cortical regions, using the fry rotation testing module in the limma R package (Wu et al., 2010).

Relationships between length-normalized log-transformed neuronal transcript abundance (transcripts per million) and global editing index were assessed with linear models, controlling for gross neuronal phenotype (excitatory or inhibitory) and library size unless otherwise stated. Gene set Gene Ontology enrichment analysis was performed on transcripts significantly associated with editing using the limma goana module (Ritchie et al., 2015), after exclusion of genes with absolute effect size estimates less than 0.01. For analyses involving ribo-depleted bulk RNAseq libraries, the proportion of brain cell types in each library was estimated using deconvolution (Hunt et al., 2019). The estimated neuronal proportion in each library was then included as a covariate when testing associations between gene expression and global RNA editing.

Data transformation and analysis was performed in R (R Core Team, 2014), using the dplyr (Wickham et al., 2015), tidyr (Wickham and Henry, 2017) and broom (Robinson et al., 2017) packages. doParallel (Weston and Microsoft Corporation, 2018) was used for parallel processing. Data were visualized using ggplot2 (Wickham, 2016), ggridges (Wilke, n.d.), ggforce (Pedersen, 2019), ggExtra (**Attali and Baker, 2016),** ggpubr (Kassambara, 2018) and UpSetR (Conway et al., 2017) packages. Scripts for reproducing results in this manuscript are available at github.com/bahlolab/brain_scRNAed.

---

**SFigure 1.**
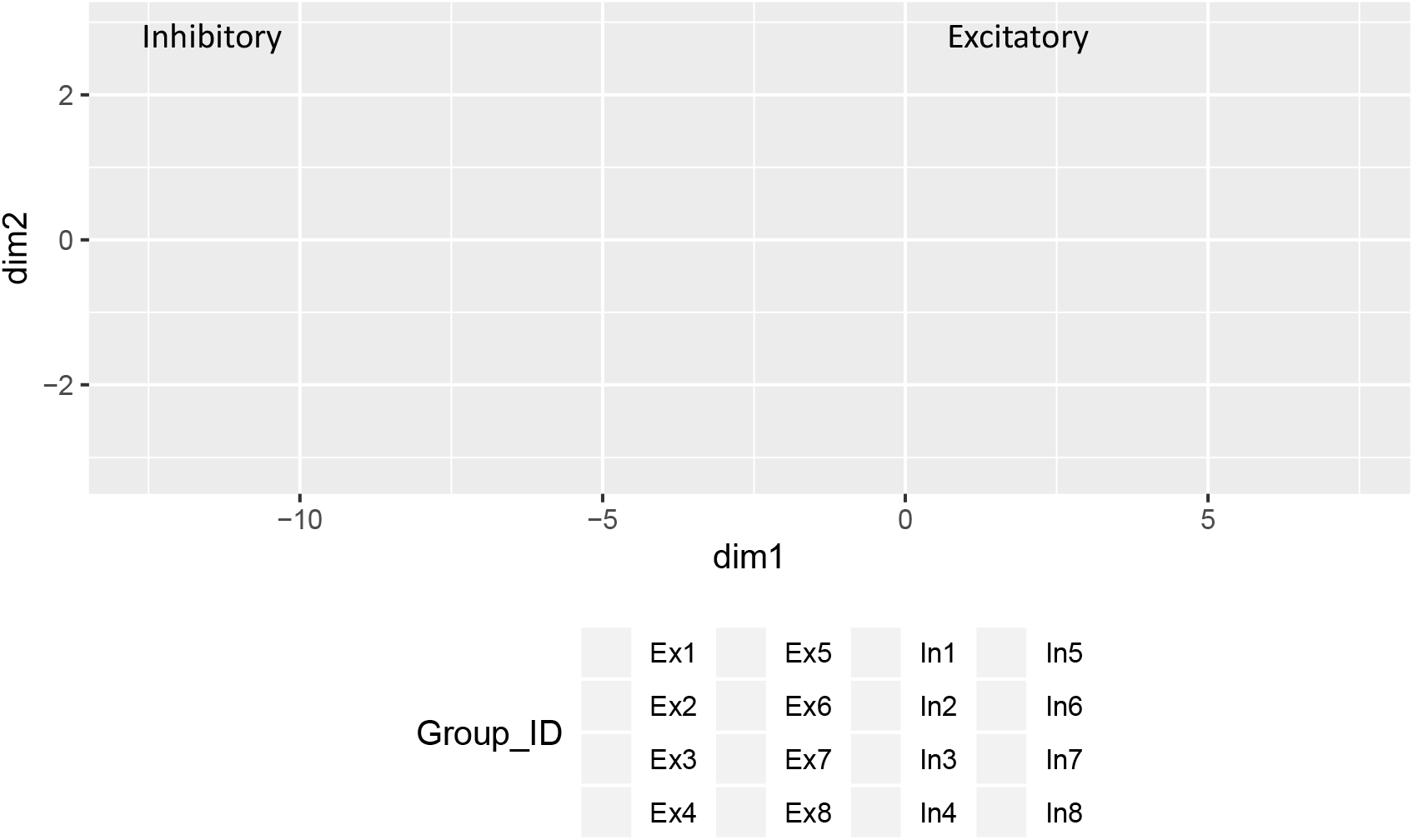
Independent clustering of 3055 high-quality neuronal nuclear transcriptomes separates inhibitory from excitatory neurons. UMAP plot displaying independent clustering of neuronal transcriptomes based on gene expression. Each point represents a neuronal nuclear transcriptome, coloured by cell type as assigned by Lake and colleagues(Lake et al., 2016). Inhibitory neurons (In) area at left, and excitatory neurons (Ex) at right. Cells with low library sizes or containing >15% mitochondrial RNA were excluded during quality control.

**SFigure 2.**
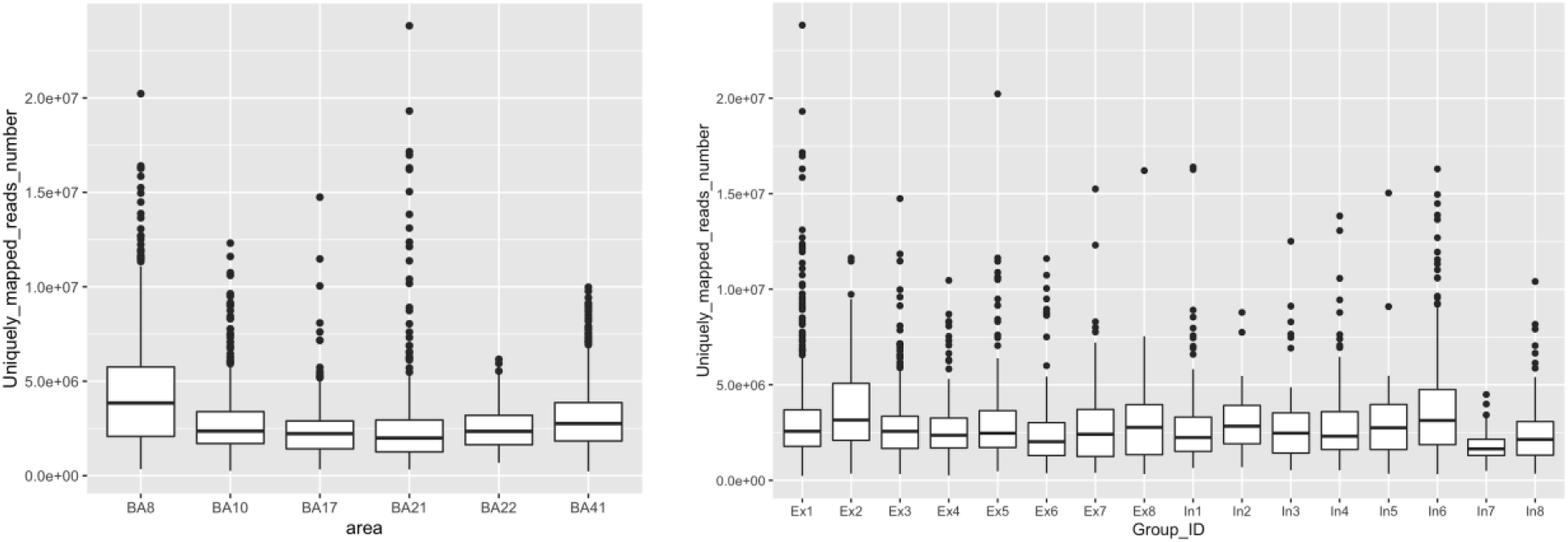
Read mapping statistics for neuronal nuclear transcriptomes, grouped by cortical area, and neuronal sub-group. The average number of uniquely mapped reads was 3 million. The mean number of multi-mapping reads was 780,535. When quantifying gene expression, a fractional assignment method was used for multi-mapping reads (see methods). BA8 showed significantly more uniquely mapped reads than other cortical regions; and neuronal sub-groups In7 and In8 had significantly lower numbers of uniquely mapped reads; whereas Ex2 and In6 had higher numbers.

**SFigure 3.**
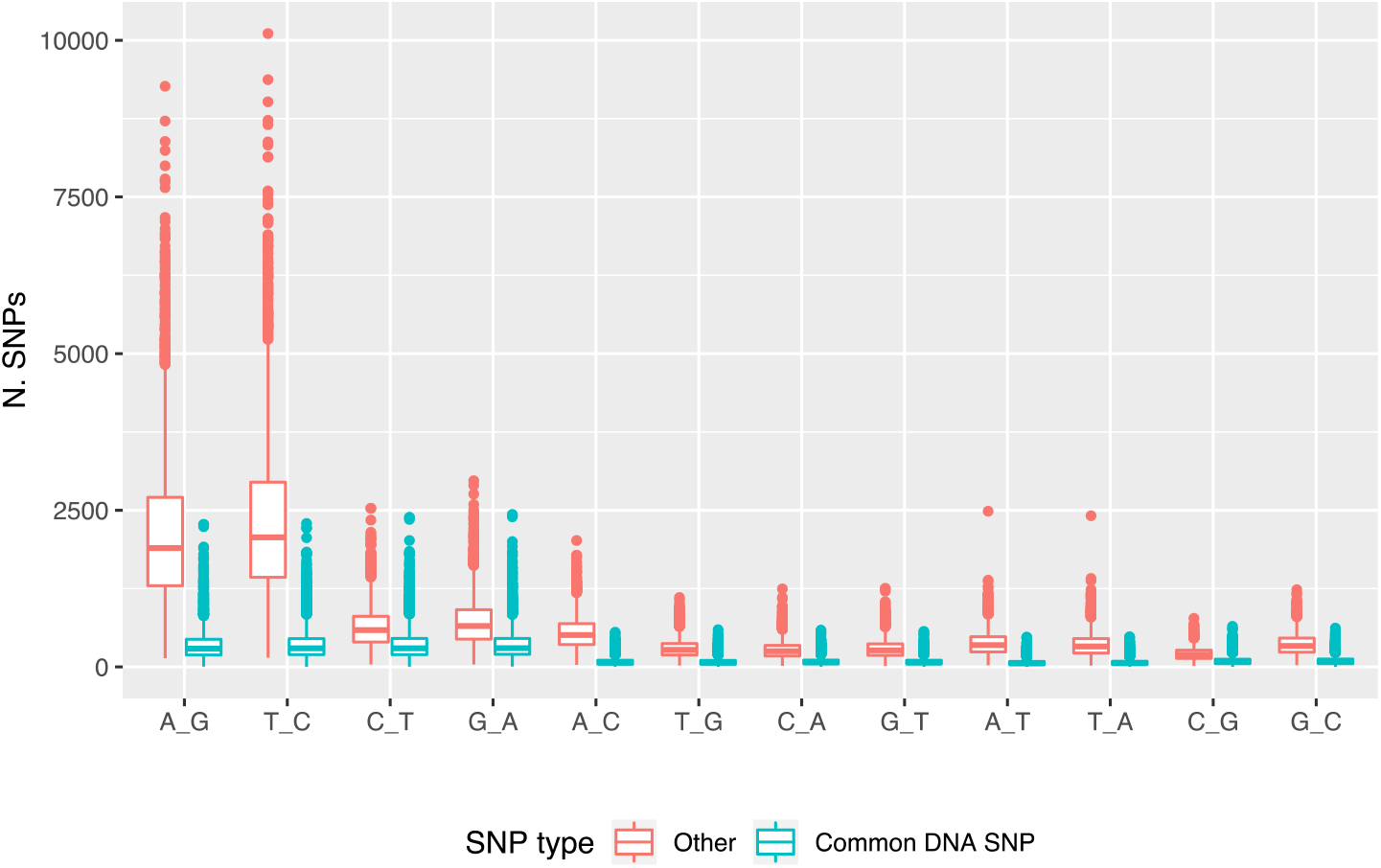
RNA editing signal is prominent in raw SNP call data. Distribution of number of SNPs (y axis) detected per nucleus in raw SNP-call data, separated by SNP type. SNP substitutions (DNA reference_RNA nucleotide) are indicated on x axis. ‘Other’ (red) SNPs encompass a strong RNA editing signal (A>G for forward DNA strand; and T>C for reverse DNA strand), distinguished from sites catalogued in the common human SNP database, dbSNP (green).

**SFigure 4.**
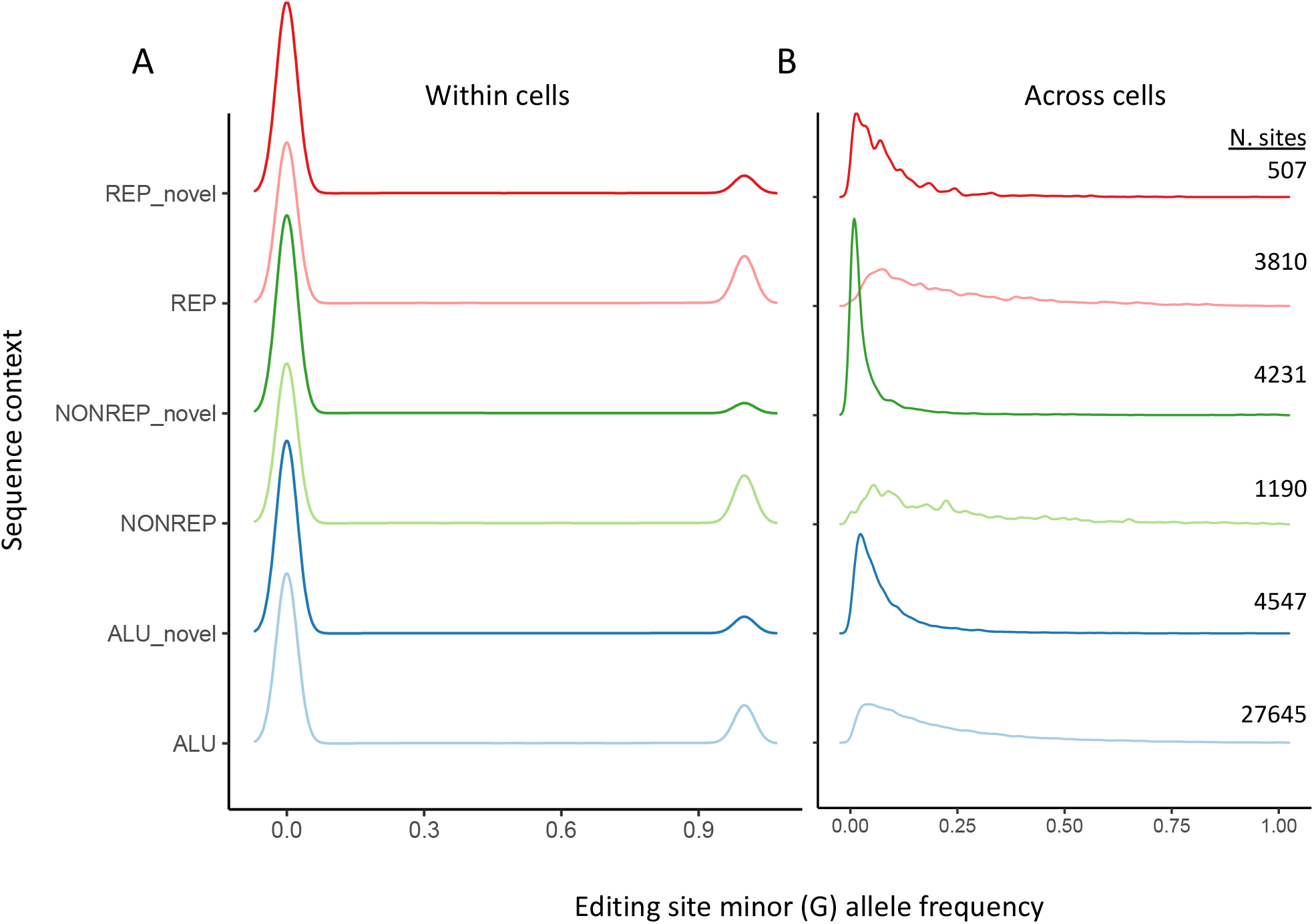
Minor allele frequency distribution for edited alleles calculated per cell (panel A) or averaged across cells (B). The sequence context for edited sites is displayed on the y axis. At the single nucleus level candidate sites are generally completely edited, or unedited. When averaged across cells, the minor allele frequency follows the positive skew reported in analyses of bulk RNA editing.

**SFigure 5.**
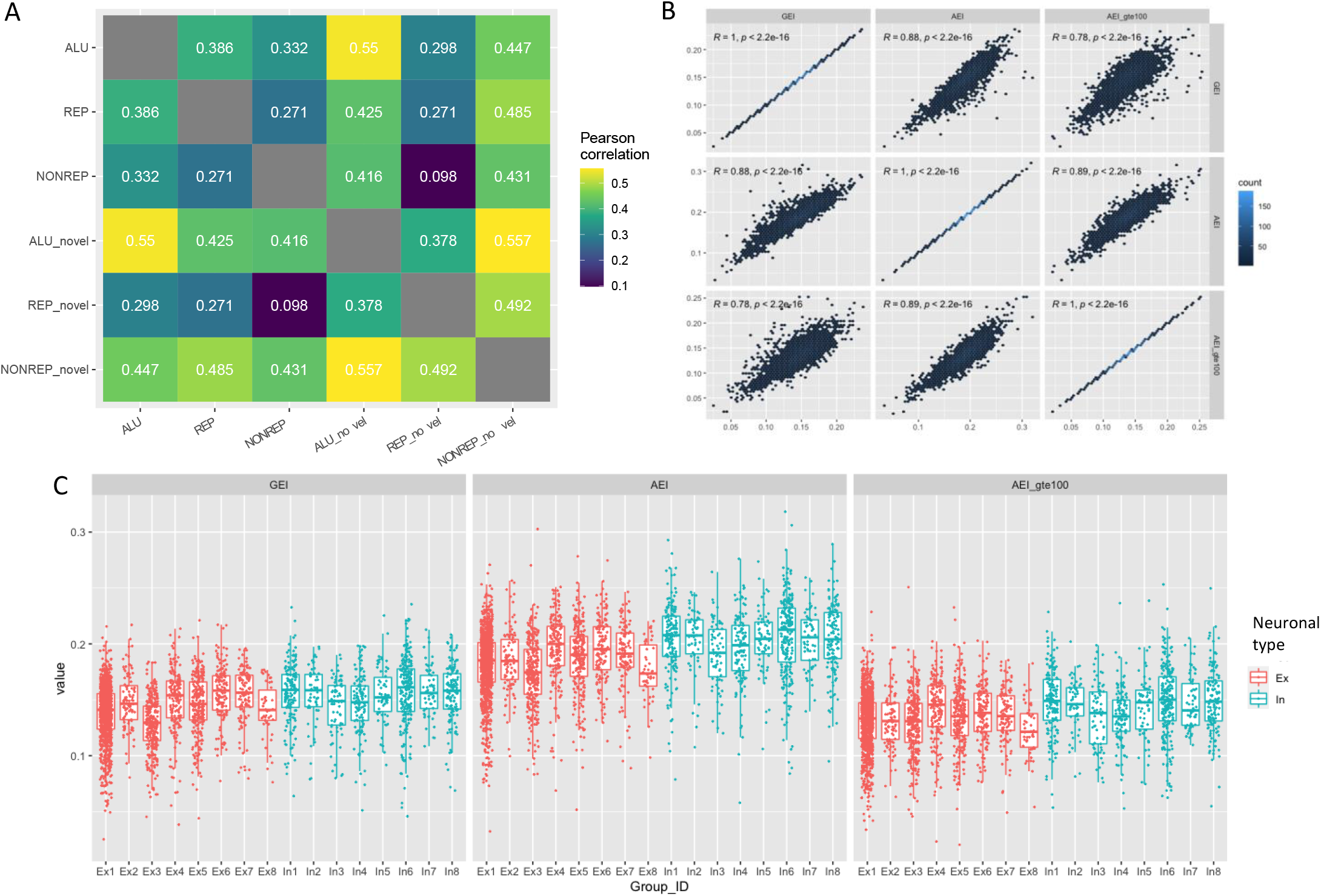
A) Correlations between log10 (number of sites per gene) for different editing sequence contexts (x and y axes). Numbers in tiles represent Pearson correlation values. Novel sites are those not previously catalogued in REDIportal, RADAR or GTEx databases, which are defined based on intersection with the Repeat Masker database (containing Alu and non-Alu repetitive regions). B) Correlations between cell-wise editing rates calculated using different sets of 41,930 high-confidence editing sites. GEI: global editing index, calculated using all transcribed candidate sites per cell; AEI: Alu editing index, calculated using all transcribed Alu sites per cell; AEI_gte100: stringent AEI, calculated using only Alu sites transcribed in at least 100 neuronal nuclei. Hexagons indicate frequency bins, coloured by number of observations per bin (‘count’). Pearson’s product-moment (R) and p value are displayed at top. C) Cell-wise editing rates (y axis) for neuronal sub-groups (x axis) calculated using the global editing index (GEI; as for Figure 3c); the Alu editing index (AEI); and the stringent AEI (AEI_gte100). Inhibitory neurons (In) show significantly greater editing than excitatory neurons (Ex) as a group irrespective of the editing index used.

**SFigure 6.**
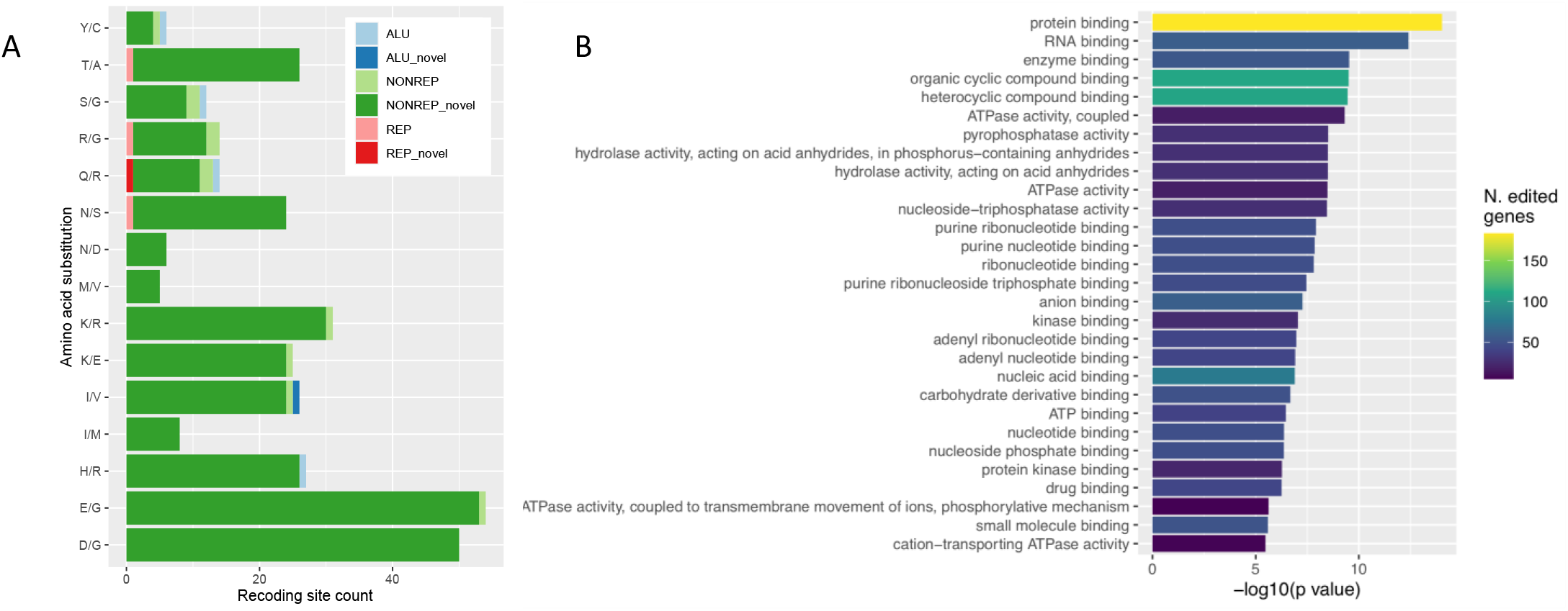
Predicted protein re-coding consequences of 316 exonic editing sites in 232 genes. A) Number of sites (x axis) encoding different amino acid substitution consequences (y axis), coloured by editing sequence context. B) Top 20 most enriched molecular function Gene Ontology terms (y axis) among 232 putative re-coding target genes. Bar length indicates enrichment significance (x axis) and fill colour indicates the number of edited genes supporting each result.

**SFigure 7.**
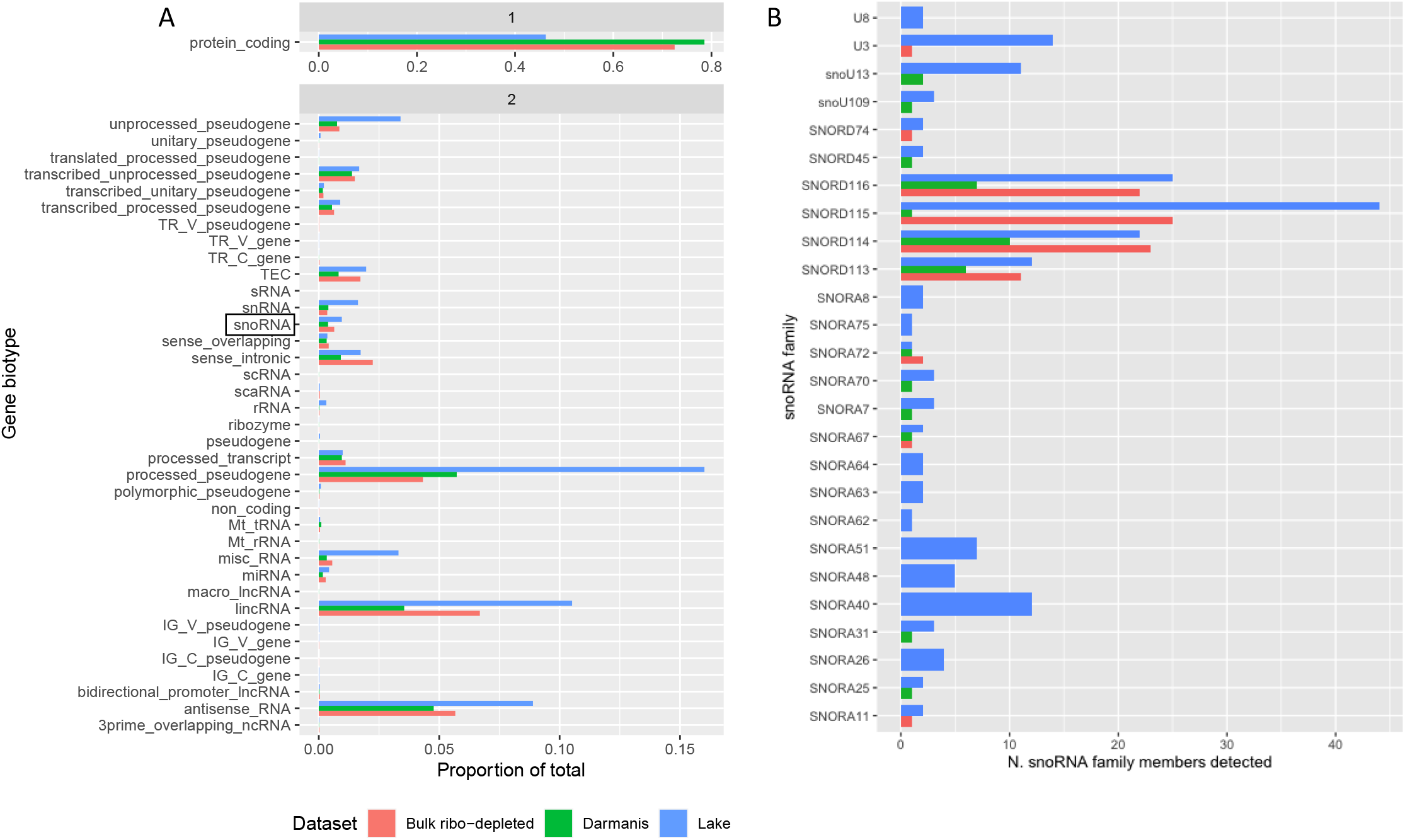
A) Proportion of gene biotypes (y axis) quantified in single nucleus RNA sequencing published by Darmanis et al. (green), Lake et al. (blue) and bulk ribo-depleted RNA sequencing (red). Note the different x axis scale for panels 1 and 2. B) Quantification of small nucleolar RNA families detected in the three data sets. Only snoRNA families with at least 5 members annotated in the human genome are displayed.

**SFigure 8.**
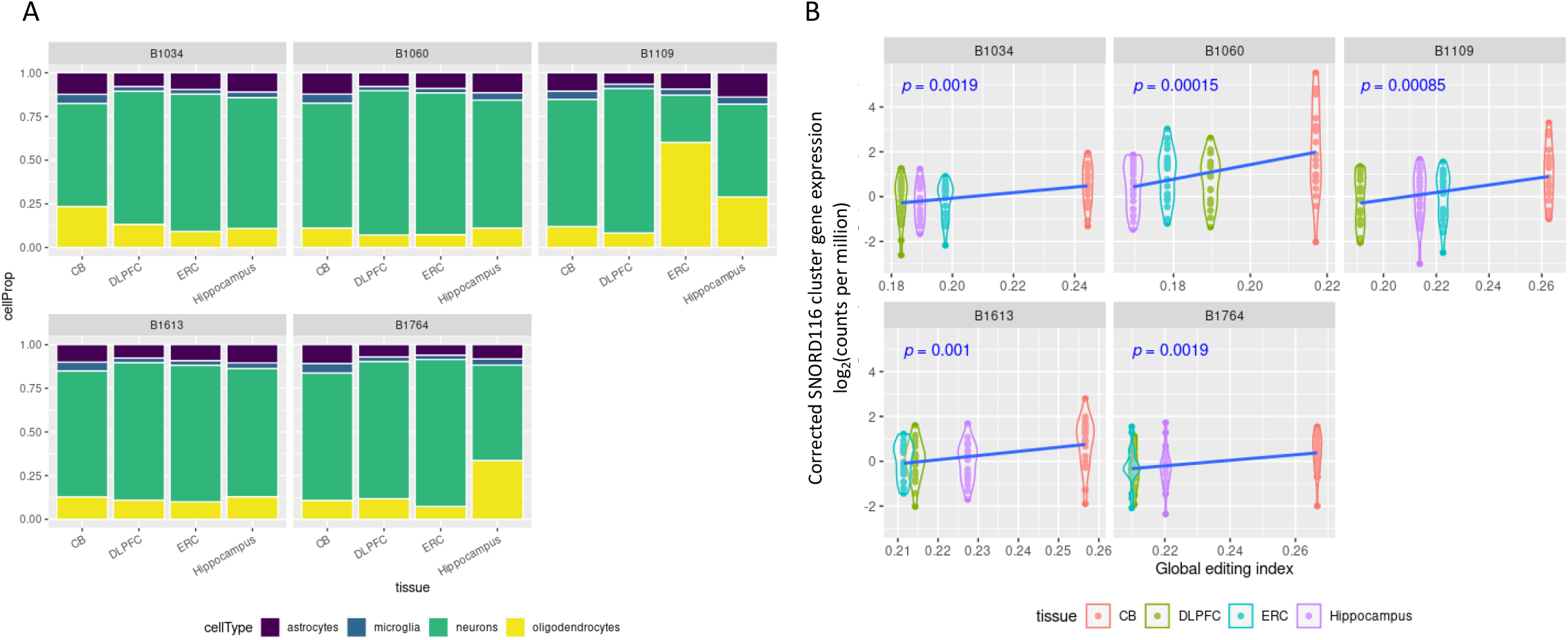
A) Cell type proportions estimated in bulk ribo-depleted RNA sequencing from four brain regions (x axis) of five adult brain donors (facets) reported by Hwang et al (Hwang et al., 2016). B) Association between expression of SNORD116 cluster genes corrected for estimated neuronal proportions (y axis), and global RNA editing (x axis) within each brain donor (facets). CB: cerebellum; DLPFC: dorsolateral prefrontal cortex; ERC: entorhinal cortex.

